# Transcriptomic responses to thermal stress and varied phosphorus conditions in *Symbiodinium kawagutii*

**DOI:** 10.1101/499210

**Authors:** Senjie Lin, Liying Yu, Huan Zhang

**Author notes:** Corresponding author (SL). These authors contributed equally to this work (Senjie Lin, Liying Yu).

## Abstract

*Symbiodinium* species are essential symbionts of tropical reef-building corals and the disruption of their symbiosis with corals as a consequence of seawater warming and other stress conditions leads to the globally widespread coral bleaching. As coral reefs live in the oligotrophic environment, *Symbiodinium* photosynthesis can also face nutrient stress. How metabolic pathways in *Symbiodinium* respond to thermal stress and phosphate depletion is poorly understood and underexplored for many species. Here we conducted RNA-seq analysis to investigate transcriptomic responses to thermal stress, phosphate deprivation and glycerol-3-phosphate (Gro3P) replacement in *S. kawagutii*. RNA-seq and bioinformatic analysis were conducted for the above-mentioned three treatments and a control. We identified 221 (2.04%) genes showing no significant differential expression among all conditions, and defined them as “core” genes of *S. kawagutii*, which mostly were in the Gene Ontology terms of catalytic activity and binding. Using algorithms edgeR and NOIseq in combination, we identified a set of differentially expressed genes (DEGs) for each treatment relative to the control. Under heat stress 357 (4.42%) DEGs were found, with predicted roles in active molecular (protein-protein/RNA/DNA) interaction, cell wall modulation and transport (including nutrients, iron, and oxygen). About as many DEGs (396, 4.73%) were identified under P deprivation while nearly double of that (671, 8.05%) were detected under Gro3P utilization; in both cases most of the DEGs were up-regulated and predicted to function in photosystem and defensome. Further KEGG pathway comparison revealed different molecular responses between phosphate deprivation and Gro3P utilization. Catalytic activity and binding seem to be two important core functions in *S. kawagutii*. The most significant transcriptional response in *S. kawagutii* to heat stress was regulation of molecular interaction, cell wall modulation, and transport of iron, oxygen, and major nutrients, suggesting that this species uses a unique mechanism to cope with heat stress, possibly conferring thermal tolerance. The greatest transcriptomic impact of phosphate deprivation and Gro3P replacement were the up-regulation of photosystem and defense. This study provides new clues about molecular mechanisms underpinning responses in *Symbiodinium* to temperature and nutrient stresses, which will generate new hypotheses and set a new framework for future investigations.

## Introduction

Dinoflagellates are known in several major roles in the ocean: important primary producers, the greatest contributors of harmful algal blooms and marine biotoxins, and essential endosymbionts of reef building corals and some other invertebrates. The coral endosymbionts come from the genus of *Symbiodinium* (1), which are classified into nine clades (A-I) based on their genetic differences despite little morphological variance (2), some of which just began to be recognized as different genera (3).

Among the nine clades of *Symbiodinium*, clade E is exclusively free-living, while the others contain endosymbionts. Coral symbionts mostly fall in clades A, B, C, D, and F (4). Our understanding on the coral-*Symbiodinium* relationship has evolved from the earlier “one coral-one genotype of *Symbiodinium*” to “one coral-one dominant genotype of *Symbiodinium*” with the discoveries that the endosymbiont assemblage contains multiple genotypes (1, 5, 6). The dominant and minor genotypes can shuffle in the process of coral bleaching (7, 8). Coral bleaching, the increasingly widespread and severe coral-degrading phenomenon, is due to the expulsion of *Symbiodinium* as a consequence of environmental stress (9). Living in the tropical oligotrophic oceanic environment, the limitation of nutrients, such as phosphorus (P) (10), in conjunction with episodes of El Nino and undergoing global warming can exacerbate coral bleaching.

It has been reported that thermal stress induces inactivation of photosystem II (PSII) in *Symbiodinium* spp. by damaging light-harvesting proteins (11). Studies have also shown that symbiosis with different genotypes of *Symbiodinium* can lead to differential susceptibility of the coral to thermal stress. For instance, *Symbiodinium* of putatively thermotolerant type D2 and the more susceptible type C3K showed markedly different gene expression profiles, especially for heat shock proteins and chloroplast membrane components within 3 days of exposure to elevated thermal exposure (12). Some studies reported depression of growth rate and photosynthesis at high temperature in clades A and B, but not in clades D and F (13). At an elevated temperature of 31 °C, a clade F showed transcriptional response in 37.01% of its 23,654 total detected unigenes, among which 2.78% exhibited ≥ 2-fold changes in expression, and these responsive genes encoded antioxidant and molecular chaperones, cellular components, and other functions (14). More studies like this on different types of *Symbiodinium* spp. can help us better understand how the symbiont community responds to environmental stress. Besides, response to nutrient stress is also important to investigate for corals and *Symbiodinium* as they typically live in the oligotrophic environment. Phosphorus is an essential nutrient for algae but its directly bioavailable form (primarily dissolved inorganic phosphate or DIP) is often limited in various parts of the global ocean (15, 16). Natural and human activities introduce dissolved organic P (DOP) into coastal waters, providing an alternative P source (17). No study has specifically addressed molecular response to P stress and replacement of DOP for phosphate in *Symbiodinium*.

We conducted a transcriptomic study on *S. kawagutii* treated with heat stress (30 °C), P deficiency, and DOP replacement. *S. kawagutii* belongs to clade F and was originally isolated from the scleractinian coral *Montipora verrucosa* in Hawaii where ambient temperature is about 25°C (18), but this specific genotype has only been occasionally found in subsequent studies (in *Pocillopora damicornis* in Heron Island (19) and in Hong Kong [Lin unpublished data], both in the Pacific). This raises a question if this is like the thermal resistant clade D genotypes that are rare under normal conditions but resistant to stress due to physiological tradeoff (20–22). A recent genome study indicated that this species possesses microRNA gene regulatory machinery that potentially targets heat shock proteins, and that the species has expanded gene repertoire of stress response (23). Besides, this species shows highly duplicated nutrient transporters including phosphate transporters and alkaline phosphatases as well as acid phosphatases that are potentially helpful for utilizing DOP. Our transcriptomic analyses to identify differentially expressed genes reported here provide insights into responses in *S. kawagutii* to both temperature and P nutrient variations.

## Results

### Overall differential gene expression profile

The RNA-seq yielded 550 to 1150 Mbp for the four culture conditions: SymkaSL1 (thermal stress at 30 °C), SymkaSL2 (control), SymkaSL3 (phosphate deprivation), and SymkaSL4 (Gro3P replacement). Approximately 60% of each dataset was successfully mapped to the genome of this species (Table 1). In total, 44.72% of the genome-predicted genes (36,850) were covered by all the transcriptomes combined. Each of the three treatments was compared with the control. To avoid inflation of DEG numbers by genes with low-expression genes, sequences that were expressed at a low level (< 10 CPM) in both two samples, or not expressed in one sample and expressed <20 CPM in the other, were excluded from the DEG analysis. This filtering resulted in a similar set of genes (~8,000, actively expressed genes or AEG) for DEG analysis for each treatment (Table 2). Of this AEG dataset, while edgeR identified 5.57% - 13.33% as DEGs, NOIseq identified 13.71% - 56.47% as DEGs. The number of genes identified as DEG by both NOIseq and edgeR was 4.42% - 8.05% of the ~8000 genes. Further functional analysis was based on this smaller set of DEGs.

**Table 1.** RNA-seq information of *S. kawagutii*.

| Sample ID | MMETSP ID | SRA ID | Condition | Clean data size (Mbp) | Mapping rate |
| --- | --- | --- | --- | --- | --- |
| SymkaSL1 | MMETSP0132 | SRR1300302 | heat stress | 660 | 59.21% |
| SymkaSL2 | MMETSP0133 | SRR1300303 | normal | 550 | 61.29% |
| SymkaSL3 | MMETSP0134 | SRR1300304 | P deprivation | 695 | 67.43% |
| SymkaSL4 | MMETSP0135 | SRR1300305 | Gro3P replacement | 1,150 | 62.34% |

\*All samples were grown in L1 medium amended natural seawater in 25 °C except listed condition (see Methods).

From the AEG set, 10,857 genes were found to be commonly expressed genes in all the four culture conditions (named AEG-C gene set) (Table 2). Of the AEG-C set, 221 (2.04%) genes showed no significant differential expression based on our strict criteria. These are considered “core” genes (CORE) in this species. The average expression of the core genes was 584 FPKM, while the highest expression was up to 13,450 FPKM. Of these 221 CORE genes, 108 (48.87%) were functionally annotatable (S1 Table). Among these annotatable CORE genes, the most highly expressed gene encodes 14-3-3 protein, and the second encodes ADP ribosylation factor. Functions of CORE genes were confirmed by GO annotation. The 84 GO annotatable genes were distributed in two subcategories of cellular component, six subcategories in molecular function and three subcategories in biological process (Fig. 1). Two of the sub-categories of molecular function were highly enriched: catalytic activity and binding. Included in the catalytic activity subcategory were oxidoreductase, hydrolase and transferase activities. There were 8 types of binding, including organic cyclic and heterocyclic compound binding, protein binding and ion binding (Fig. 1).

**Table 2.** Number of DEGs under different conditions identified by NOIseq and edgeR.

| Group | AEG | NOIseq | edgeR | NOIseq+edgeR |
| --- | --- | --- | --- | --- |
| HS | 8,081 | 1,108 (13.71%) | 450 (5.57%) | 357 (4.42%, 249 ↑ +108 ↓ ) |
| -P | 8,364 | 1,535 (18.35%) | 557 (6.66%) | 396 (4.73%, 332 ↑ +64 ↓ ) |
| OP | 8,335 | 4,707 (56.47%) | 1,111 (13.33%) | 671 (8.05%, 580 ↑ + 91 ↓ ) |
| Total union | 10,857 | 5,397 | 1,601 | 1,091 |

\* AEG: actively expressed genes (average CPM $\geq 10$ ). HS: heat stress; -P: P deprivation; OP: organophosphate.

**Fig. 1.**
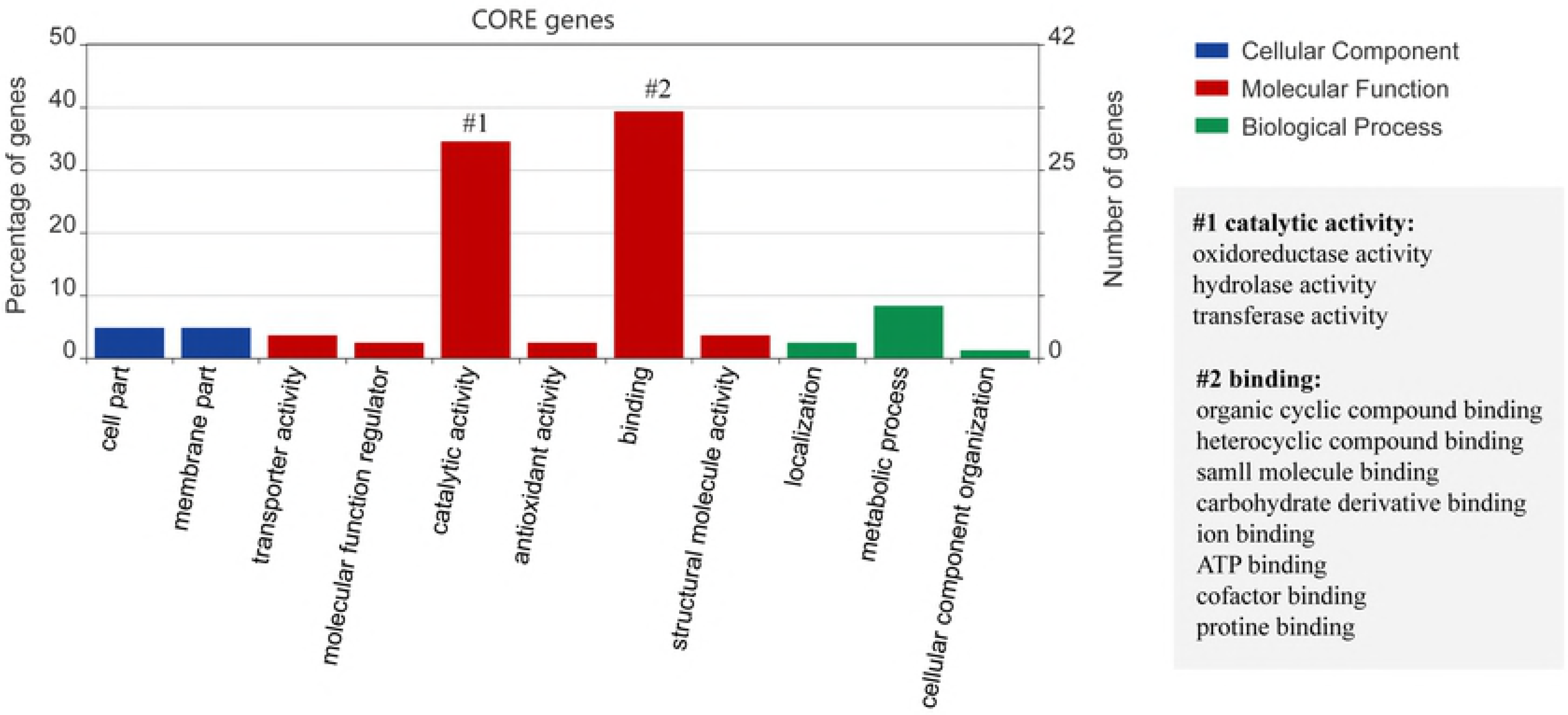
Core genes GO annotation category. Core genes were defined as genes commonly detected under all four conditions: control, heat stress, P deprivation, and DOP replacement, and showed expression of ≥ 50th percentile average FPKM and low CV of ≤ 0.1.

### Functional distribution of DEGs responding to heat stress

Under thermal stress, *S. kawagutii* showed 357 (4.42%) DEGs, with over two-thirds being up-regulated and nearly one-third being down-regulated (Table 2). Of these DEGs, 171 genes (47.90%) were functionally annotatable (S2 Table). As expected, expression of heat shock proteins (HSPs) including HSP40, HSP70, and HSP90 and chaperonin Cpn60 was strongly elevated during heat stress. Furthermore, the abiotic stress-induced glutathione s-transferase was transcriptionally promoted by thermal stress. A phytoglobin (Skav228962), a plant counterpart of animal hemoglobin involved in binding and/or transporting oxygen (24), previously suggested as a stress biomarker (25), was also up-regulated. Reversely, the conjugative protein genes including ATP aldo/keto reductase family and sulfotransferase were depressed. Photosystem I reaction center subunit IV and two iron permease (FTR1) genes involved in iron uptake were also markedly suppressed (by 5-10 folds). In transport activities, amino acid transporter, formate/nitrite transporter, ion transport protein and p-type ATPase transporter were highly up-regulated. In contrast, choline transporter-like protein, nucleotide-sugar transporter and ATP binding cassette (ABC) transporters were down-regulated. In signal transduction process, most of genes encoding protein kinases, ATPase and polycystin 2 showed elevated expression under heat stress. Meanwhile, GO enrichment analysis indicated that three GO terms were significantly enriched (Fig. 2). Carbohydrate metabolic process was one of them, which consisted of a down-regulated alpha-D-phosphohexomutase, three up-regulated glycoside hydrolases and an up-regulated fructose-bisphosphate aldolase. Iron binding activity was potentially promoted as inositol oxygenases and cytochrome c/P450 genes were significantly up-regulated. In addition, cytoplasm was enriched with markedly up-regulated inositol oxygenases, inorganic pyrophosphatase and protein CfxQ. Interestingly, a protocaderin fat gene was highly expressed and induced by heat stress, so was cytokinin riboside 5’-monophosphate phosphoribohydrolase (Fig. 2).

**Fig. 2.**
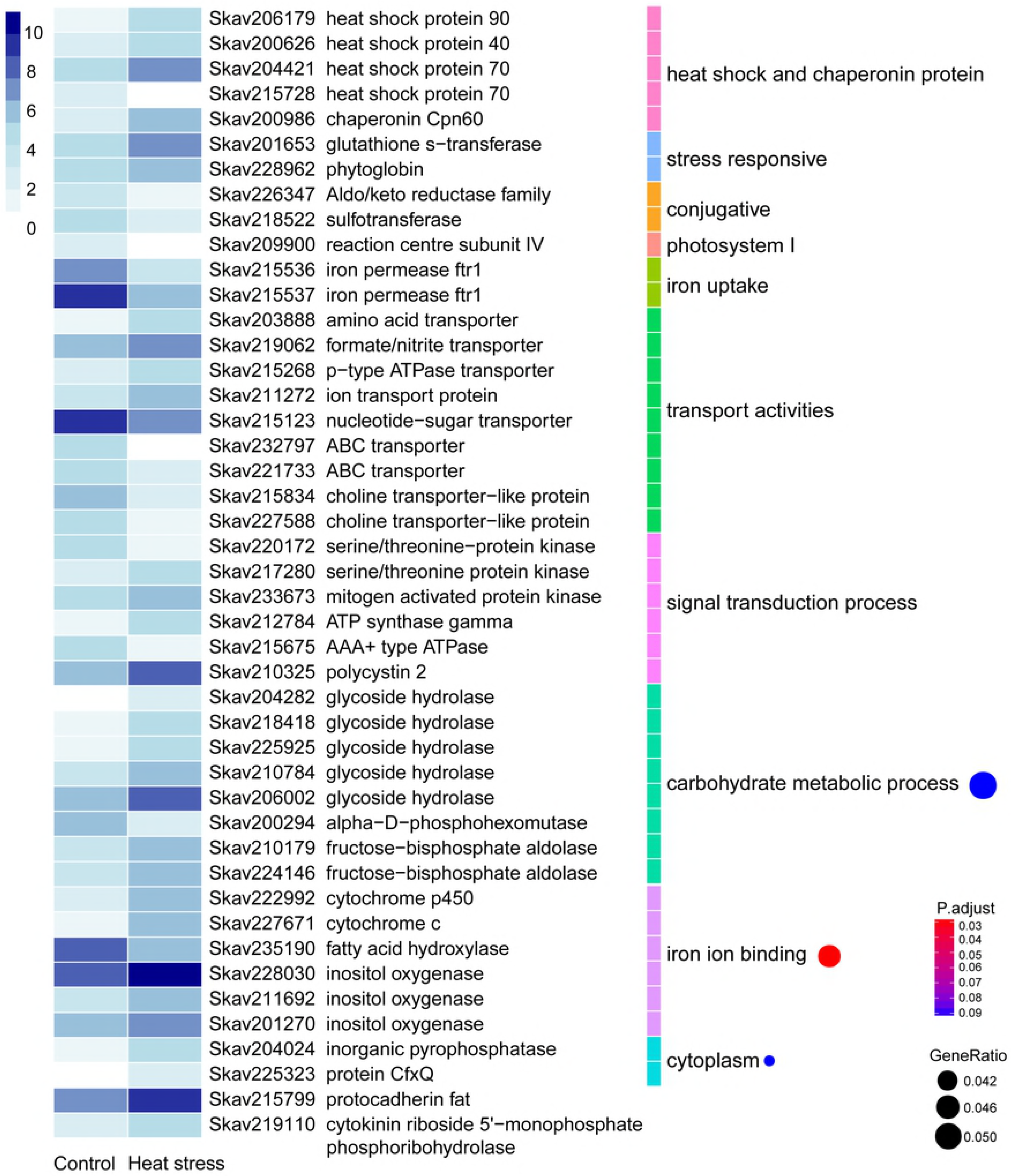
Differential gene expression in response to heat stress (30 °C). The heatmap color strength represents log2-transformed gene expression level estimated as <u>F</u>ragments <u>P</u>er <u>K</u>ilo-base of exon model per <u>M</u>illion mapped fragments (FPKM), from dark blue (highest), light blue, to white (lowest). Each color bar on the middle right marks a functional category. Each dot on the right marks a category based on GO enrichment (p-value cutoff = 0.1). The dot size represents enriched DEGs count. The color strength represents the P-value. Control: SymkaSL2 grown in L1 medium at 25 °C. HS: SymkaSL1 was grown in L1 medium at 30 °C.

Besides, duplicated gene families as reported in the *S. kawagutii* genome (23) were up-regulated, including most of ankyrin repeat (AR)-, F-box and FNIP repeat- and pentatricopeptide repeat (PPR)- containing proteins, and EF-hand domain-containing calcium-binding proteins, Zinc finger (ZnF) proteins, glycoside hydrolases, regulator of chromosome condensation (RCC1), and glycosyl transferases (S2 Table).

### Functional distribution of DEGs responding to P stress

DEG analysis revealed that 396 genes (4.73%) were responsive to P stress (Table 2). More than 80% of these DEGs exhibited significantly higher expression levels than the control whereas only 64 showed down-regulation. Of the 396 DEGs, 231 (58.3%) were functionally annotatable (S3 Table). Among the up-regulated were genes coding for proteins involved in phosphate exchange between chloroplast and cytoplasm, triose phosphate/phosphate translocator (TPT) (26), and a score of phosphatases potentially involved in DOP metabolism and utilization, such as alkaline phosphatase (AP), phosphoserine phosphatase, phosphoglycolate phosphatase, protein phosphatase 2C, and metal-dependent phosphohydrolase. Differential gene expression was richly represented in photosynthetic apparatus under the influence of P stress: photosystem II (PSII) light harvesting complex proteins and chlorophyll a-c binding protein, photosystem I (PSI) reaction center subunit IV and PsaDs, as well as electron-transfer proteins in photosynthetic electron transfer chain of flavodoxin and ferredoxin. Under P stress, a total of 11 differentially expressed PSII chlorophyll a-c binding protein genes were up-regulated, and two light harvesting complex protein genes were down-regulated. Eight of the 11 PSII chlorophyll a-c binding protein genes were enriched in GO term of photosynthesis light harvesting (Fig. 3). The results were consistent with changes in gene expression of *Prymnesium parvum* induced by nitrogen and phosphorus limitation (27). Two PSI PsaD genes were up-regulated but the PSI reaction center subunit IV decreased to an undetectable level in the P-deprived cultures. GO enrichment showed that these three genes were concentrated in photosystem I reaction center (Fig. 3). Interestingly, the highly expressed flavodoxin was decreased while ferredoxin was enhanced under P stress; flavodoxin is known to be induced under iron limitation to replace its iron-containing functional equivalent ferredoxin in diatoms and some other algae (28–31). P plays an important role in the production of ATP, NADH and NADPH; thus, energy related genes of six DEGs were explored. They included three ATPase genes, a p-type ATPase transporter, glyceraldehyde 3-phosphate dehydrogenase and geranylgeranyl diphosphate reductase. Among them five were up-regulated while ATPase subunit C was down-regulated under P stress. Besides, oxidoreductase activity and heme binding terms were enriched, mostly comprised of up-regulated haem peroxidase, globin, cytochrome b5 and P450 (Fig. 3).

**Fig. 3.**
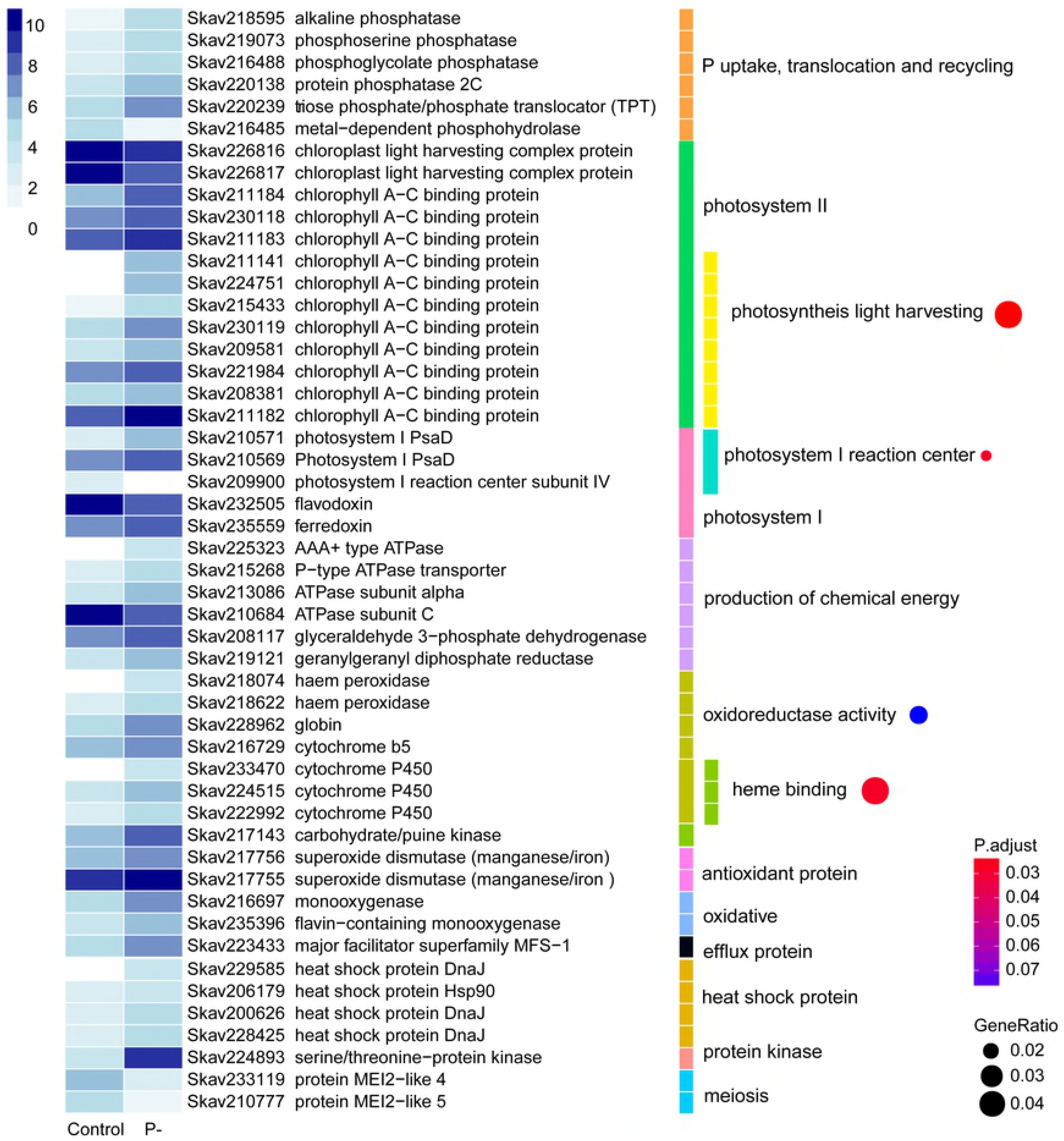
Differential gene expression in response to phosphate deprivation. The heatmap color strength represents log2-transformed gene expression by <u>F</u>ragments <u>P</u>er <u>K</u>ilo-base of exon model per <u>M</u>illion mapped fragments (FPKM), from dark blue (highest), light blue, to white (lowest). Each color bar on the middle marks a functional category. Each dot on the right marks a category based on GO enrichment (p-value cutoff = 0.1). The dot size represents enriched DEGs count. The color strength represents the P-value. Control: SymkaSL2 grown in L1 medium at 25 °C. P-: SymkaSL3 was grown in L1 medium with depleted DIP at 25 °C.

Under P deprived condition, several stress responsive genes were observed (Fig. 3). The antioxidant protein genes encoding superoxide dismutase were highly expressed and induced. Two oxidative genes, flavin-containing monooxygenases and the major facilitator superfamily MFS-1 were induced under P-stress condition. Also, heat shock proteins DnaJ and Hsp90 were differentially expressed. Several genes of protein kinases were up-regulated. Among these, a serine/threonine-protein kinase (Skav224893), showing 70% sequence similarity to rice PSTOL1 whose overexpression significantly enhances rice productivity under low phosphorus conditions (32), was expressed at a nearly 30 folds higher level under P stress. Gene homologs of MEI2, a meiosis associated gene in yeast, were sharply down-regulated. Some expanded gene families reported in the genome were also detected as DEGs, such as P-stress responsive up-regulated AR repeat genes (S3 Table).

### Functional distribution of DEGs responding to DOP replacement

When the preferable DIP in the growth medium was replaced by the same molar concentration of glycerophosphate (Gro3P), the transcriptionally responsive DEG set in *S. kawagutii* changed. In total, 671 (8.05%) DEGs were identified, which composed 580 up- and 91 down-regulated genes (Table 2). Of all the 671 DEGs, 347 matched a functionally annotated gene in the databank (S4 Table). A total of 8 genes were related to P utilization and exhibited higher expression levels under Gro3P as P-source than DIP (Fig. 4). Among them were two genes annotated as acid phosphatase that are believed to facilitate utilization DOP under DIP deficiency in plants (33). Photosynthesis related genes were also markedly regulated by DOP. These included 17 DEGs of PsaD and PsaL in PSI, ferredoxin and cytochrome c6 in the photosynthetic electron transfer chain, PSII cytochrome b559 and chlorophyll a-c binding protein, as well as chlorophyll synthesis enzyme protochlorophyllide reductase. All PS related genes were promoted except cytochrome b559 that was down-regulated.

**Fig. 4.**
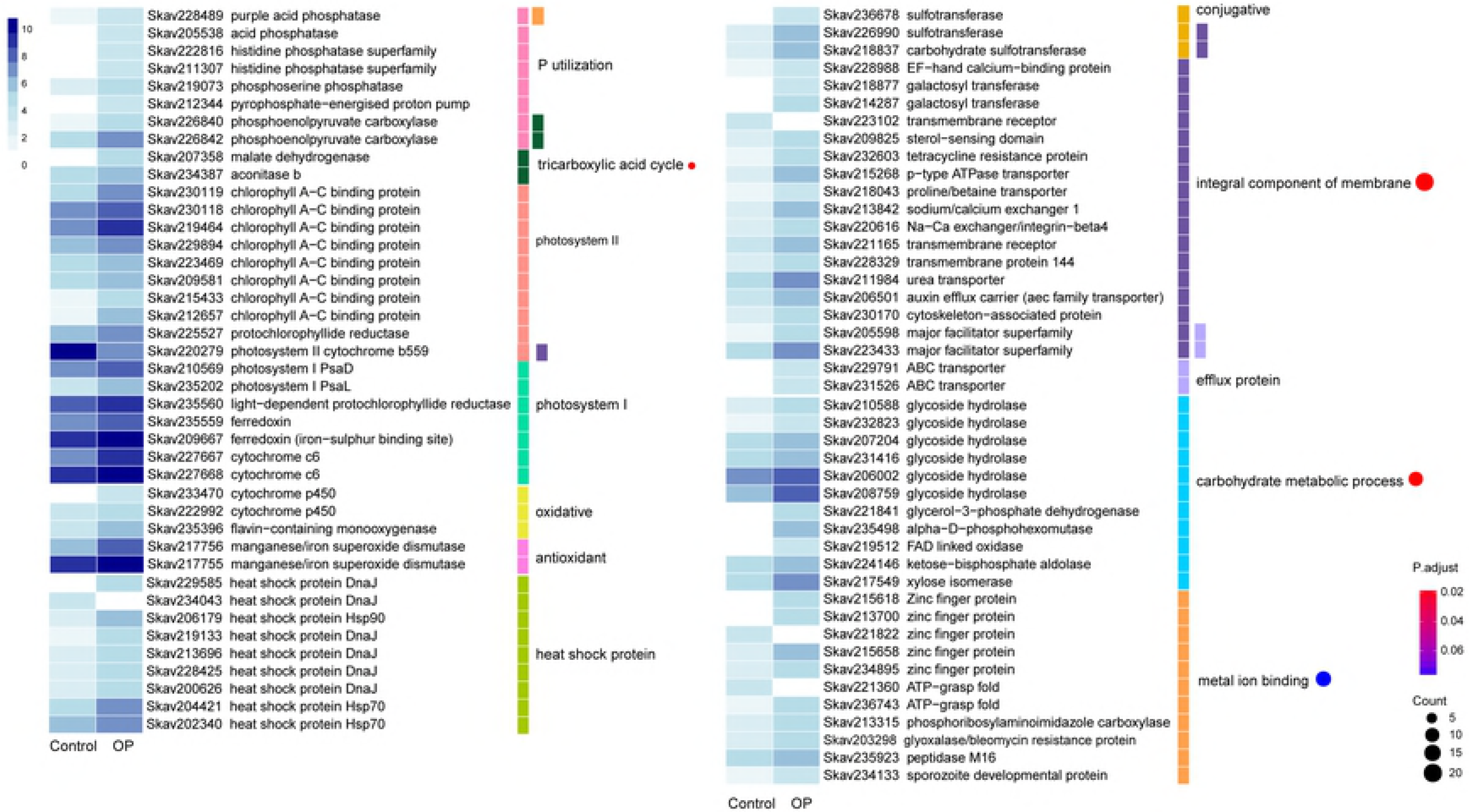
Differential gene expression in response to organophosphate. The heatmap color strength represents log2-transformed gene expression by <u>F</u>ragments <u>P</u>er <u>K</u>ilo-base of exon model per <u>M</u>illion mapped fragments (FPKM), from dark blue (highest), light blue, to white (lowest). Each color bar on the middle right marks a functional category. Each dot marks a category based on GO enrichment (p-value cutoff = 0.1). The dot size represents enriched DEGs count. The color strength represents the P-value. Control: SymkaSL2 was grown in L1 at 25 °C. OP: SymkaSL4 was grown in L1 medium with DIP replaced by glycerophosphate at 25 °C.

The defensome genes (34) were strongly induced by organophosphate replacement (Fig. 4). These included the efflux pump ABC transporter and major facilitator superfamily, oxidative proteins cytochrome P450 and flavin-containing monooxygenase, conjugative enzyme sulfotransferase, and antioxidant proteins manganese/iron superoxide dismutase. Nine heat shock proteins including DnaJ, Hsp70 and Hsp90 were also observed to be DOP-responsive. In addition, GO enrichment analysis showed that four terms were significantly impacted by the DOP replacement, including 22 DEGs specifying integral components of membrane, 17 DEGs regulating metal ion binding, 4 DEGs involved in tricarboxylic acid cycle (TCA cycle), and 12 DEGs associated with carbohydrate metabolic process (Fig. 4). Specially, two glyceraldehyde 3-phosphate dehydrogenase genes were greatly up-regulated in the DOP treatment (S4 Table). The expanded gene families identified as P-stress inducible above were also significantly regulated under DOP (S4 Table).

### Comparison of DEGs between P stress and DOP replacement

The functional diversity of genes regulated by P stress was almost the same as that by DOP replacement, but the number of responsive gene families was different between the two treatments. There were 207 DEGs commonly responsive to P stress and DOP replacement in comparison to the control. Besides, 189 and 464 DEGs were unique to P stress and DOP replacement, respectively (Fig. 5A). Overall many photosynthesis genes were affected by both P conditions, although the impacted components were different. To uncover gene expression differences in *S. kawagutii* between P stress and DOP replacement, KEGG pathway enrichment (pvalue cutoff = 0.5) comparison on DEGs was performed. As Fig. 5B shows, P stress and DOP replacement induced 88 and 137 orthologs of DEGs, respectively, involving 11 shared pathways in both groups. The common pathways included dominant lysosome, ubiquitin mediated proteolysis, oocyte meiosis, protein processing in endoplasmic reticulum, photosynthesis, TCA cycle, among others. For the oocyte meiosis pathway, genes annotated as best match in InterPro databases of S-phase kinase-associated protein, serine/threonine-protein kinase, 14-3-3 proteins, and Poly(ADP-ribose) polymerase were up-regulated under both P treatments. Pathways specifically enriched under P stress included cell cycle, cytochrome P450, TGF-beta signaling pathway, pentose and glucoronate interconversions, antenna proteins, arginine and proline metabolism, plant hormone signal transduction, ascorbate and aldarate metabolism, hippo signaling pathway and apoptosis. In contrast, DOP replacement uniquely impacted 22 pathways, including carbon metabolism and fixation, RNA degradation, and longevity regulating pathway (Fig. 5B).

**Fig. 5.**
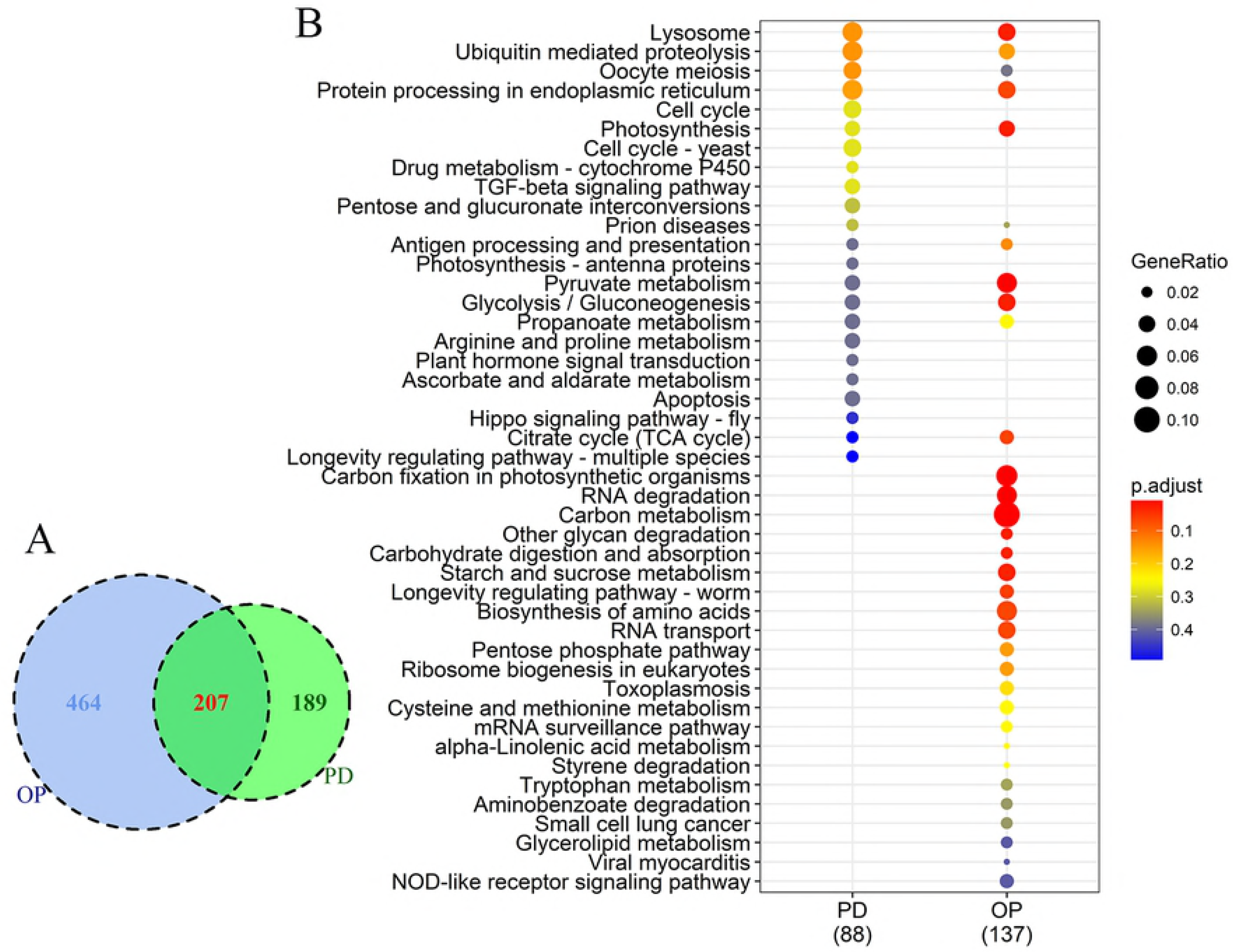
Comparison of differentially expressed genes between P-deprived (PD) and organophosphate (OP) conditions. A) Venn diagram showing numbers of DEGs common in both or unique to each of the two P conditions. B) KEGG pathway enrichment; dot size represents enriched DEGs count; color strength represents the P-value (from lowest in red to highest in blue).

## Discussion

Our transcriptome sequencing had a limited depth coverage and lacked biological replicates due to high sequencing cost before 2011 when this project was conducted under the Marine Microbial Eukaryote Transcriptome Sequencing Program (MMETSP) (35). To maximize the reliability of the results given the limitation, we took an as robust as possible and conservative approach in data analysis. First of all, we used edgeR (medium BCV=0.2) (36)and NOIseq (37) in combination to identify differentially expressed genes (DEGs). Furthermore, recognizing that due to the low depth coverage we likely missed low-expression genes, our interpretation of data will be framed on highly expressed genes. In addition, sequence reads that matched multiple genes and genes with low read counts (<10 count per million)(38), which is attributed to either short genes expressed at low levels or genes with small fold changes (39, 40), was discarded to reduce false-positive DEG calls. Consequently, the detected DEGs in this study were supposed to be highly expressed and significantly regulated genes in *S. kawagutii* under the conditions examined in this study.

Given our conservative way of identifying DEGs, it is no surprise that a smaller set of DEGs (357) responding to heat stress was identified in our study than that reported previously by Gierz et al. (14), in which 1,776 DEGs with ≥ 2-fold change in expression were found in *Symbiodimium* spp. exposed to 31 °C (compared to 24.5 °C as control), accounting for 7.51% of the transcriptome (~23,654 unique genes). However, our smaller DEGs datasets were consistent with the previous larger datasets in revealing stress responsive heat shock and chaperonin proteins, ubiquitin proteasome, and alteration in carbohydrate metabolic process (Table 3). Our DEG analysis in addition revealed a diverse set of genes that were transcriptionally regulated under heat stress, including high-affinity iron permease and iron binding molecules, oxygen transporter (phytoglobin), and genes specifying nutrient transport activities and cell features.

The results from the current study, including the DEGs and the commonly expressed genes identified, provide a new perspective and a number of previously unsuspected processes or molecular functions involved in stress response of *Symbiodinium*. This study also is the first to explore transcriptomic responses to P deprivation and replacement of DIP with DOP. The results also raise many new questions to be addressed in future research, which have high potential to lead to new insights into triggers and processes of coral bleaching and other stress symptoms. These constitute a valuable genomic resource for further inquiries into mechanisms by which *Symbiodinium* spp. (and corresponding coral host) respond to and resist environmental changes and stress.

**Table 3.**
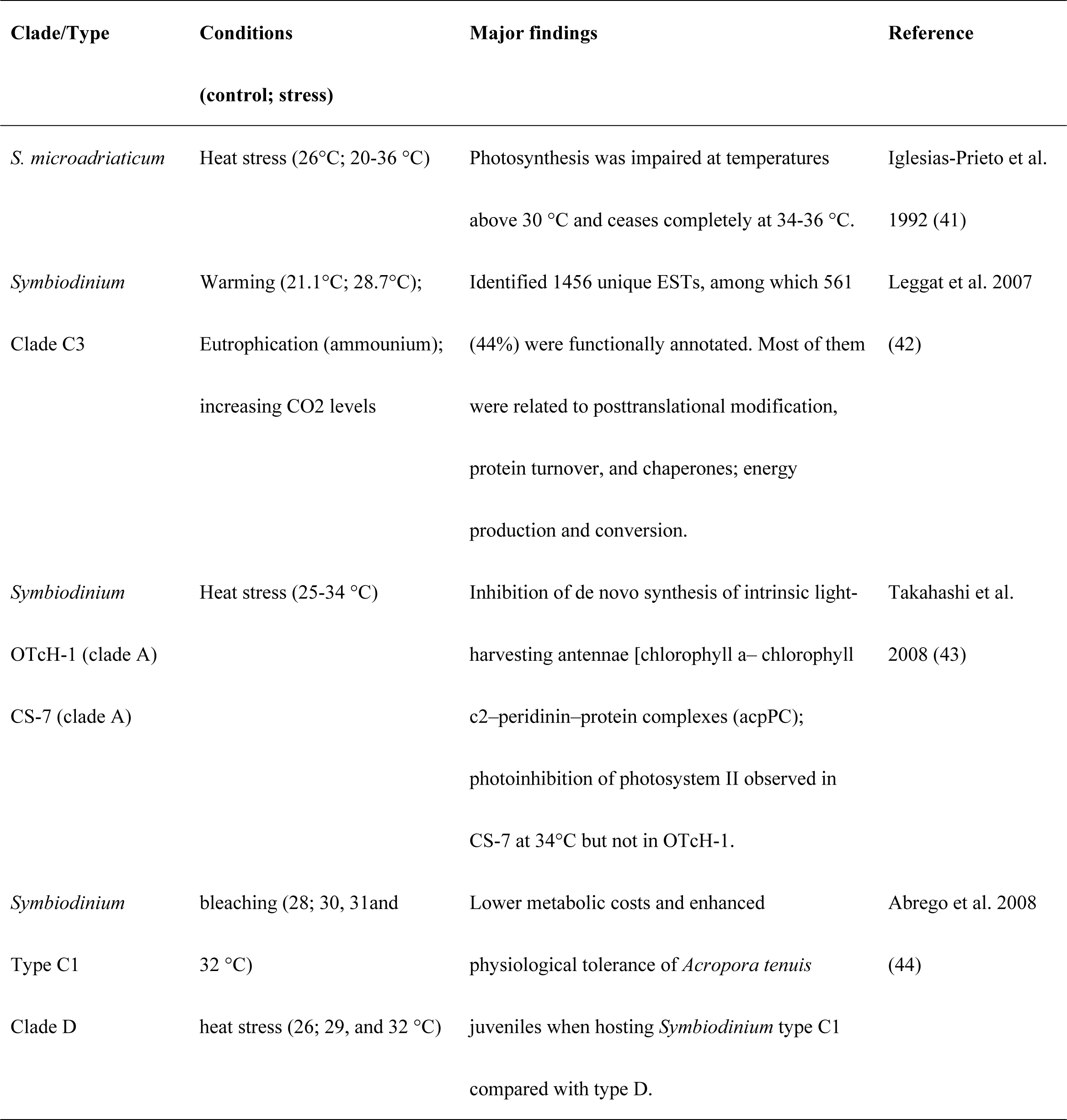
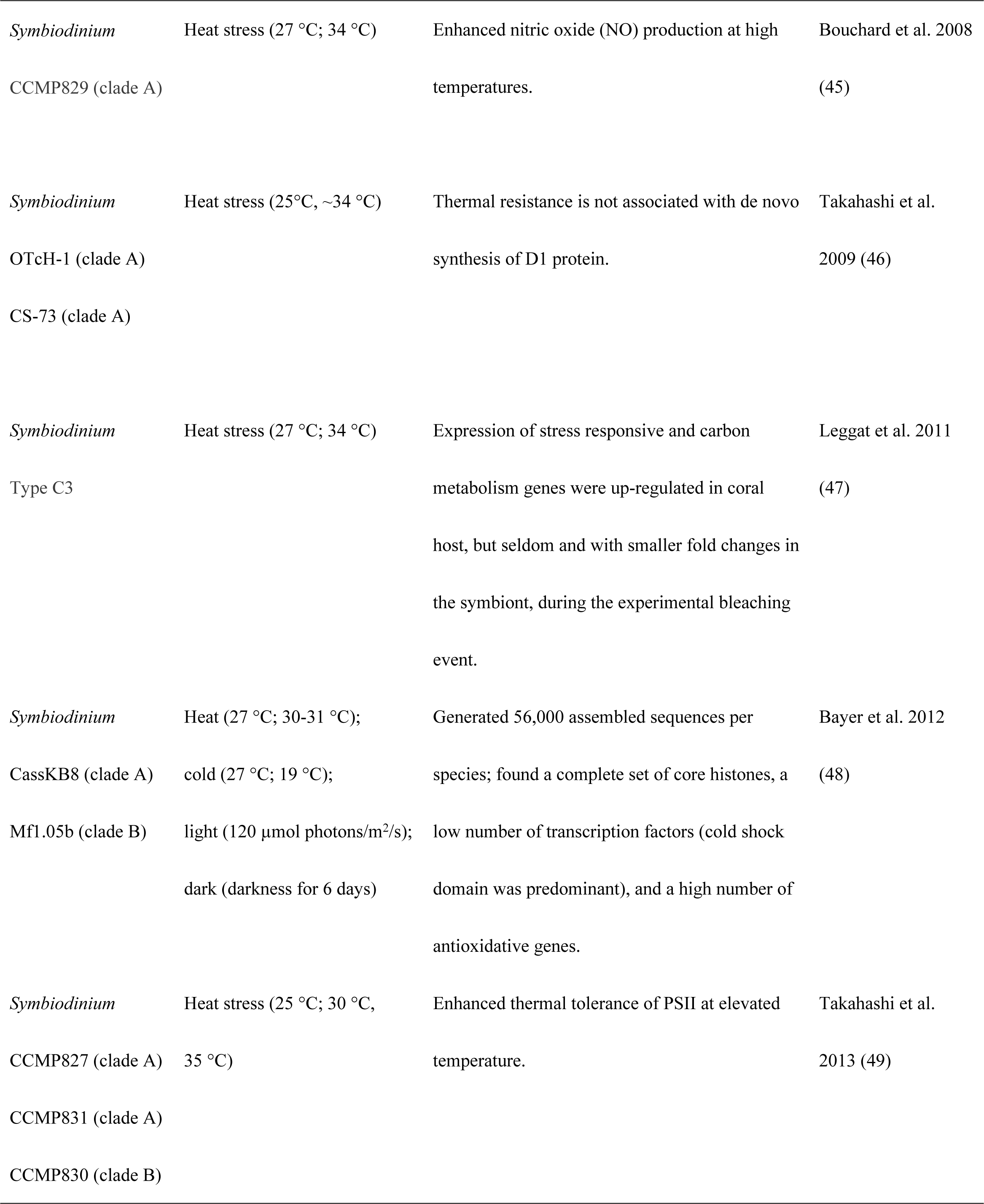
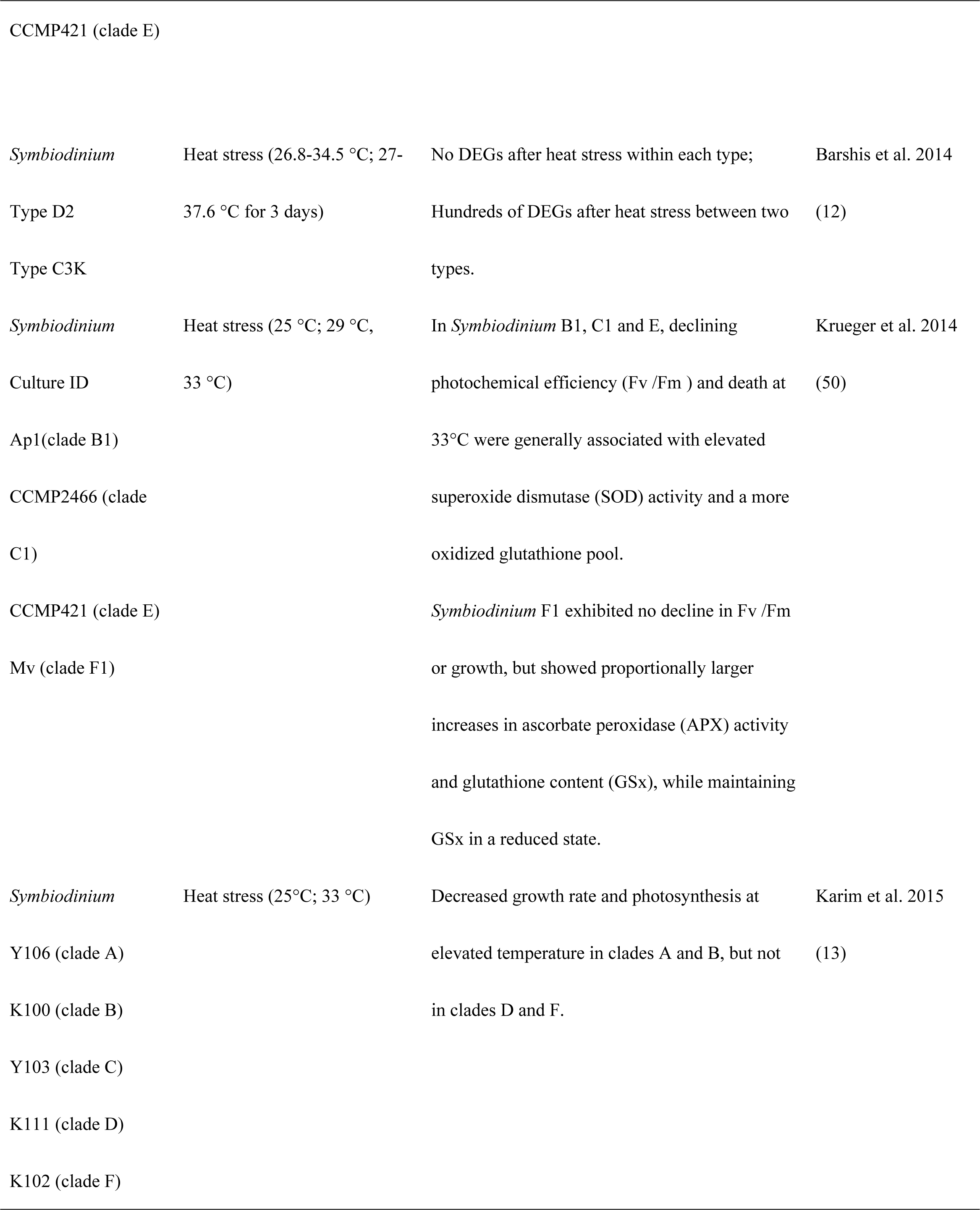
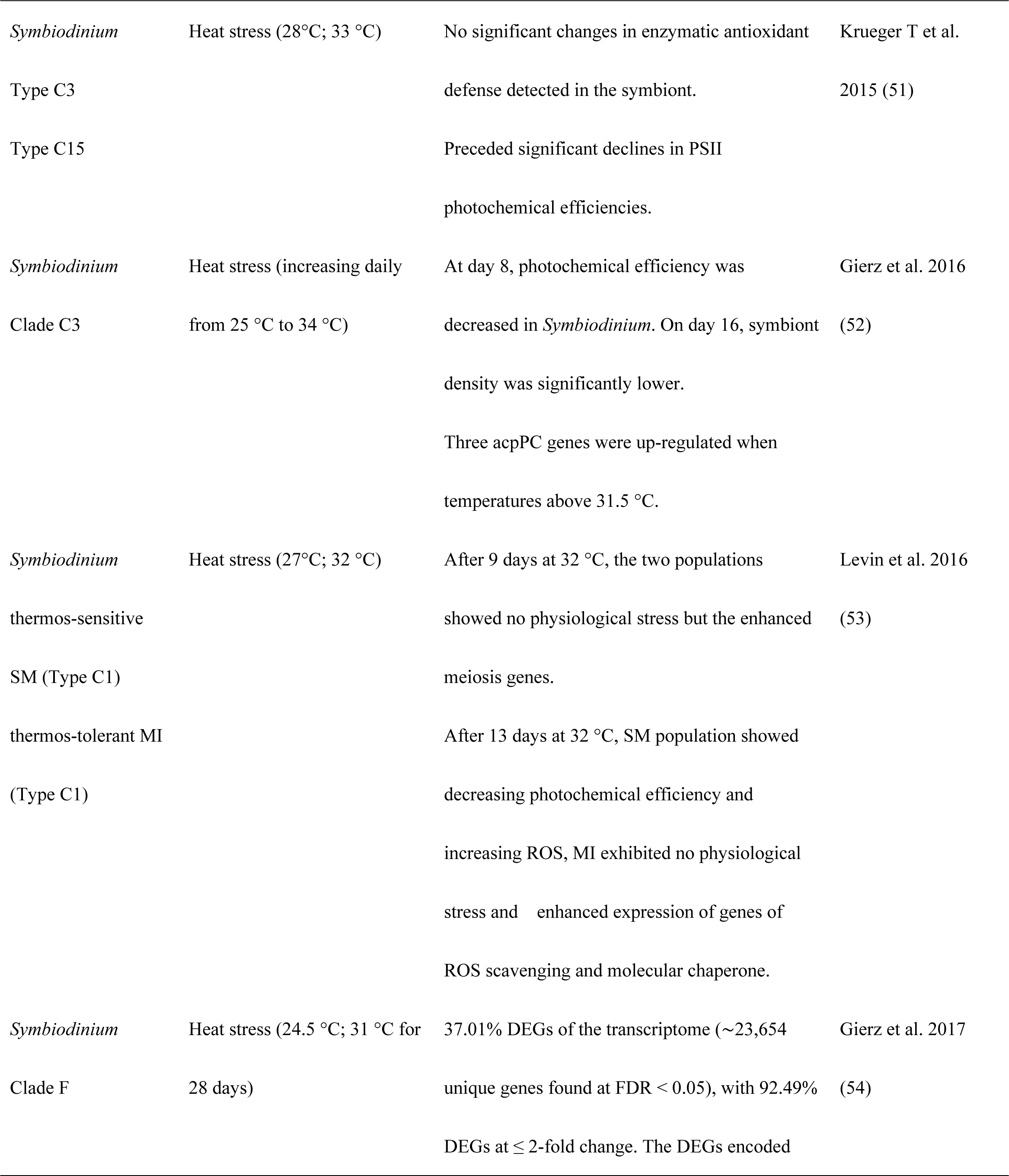
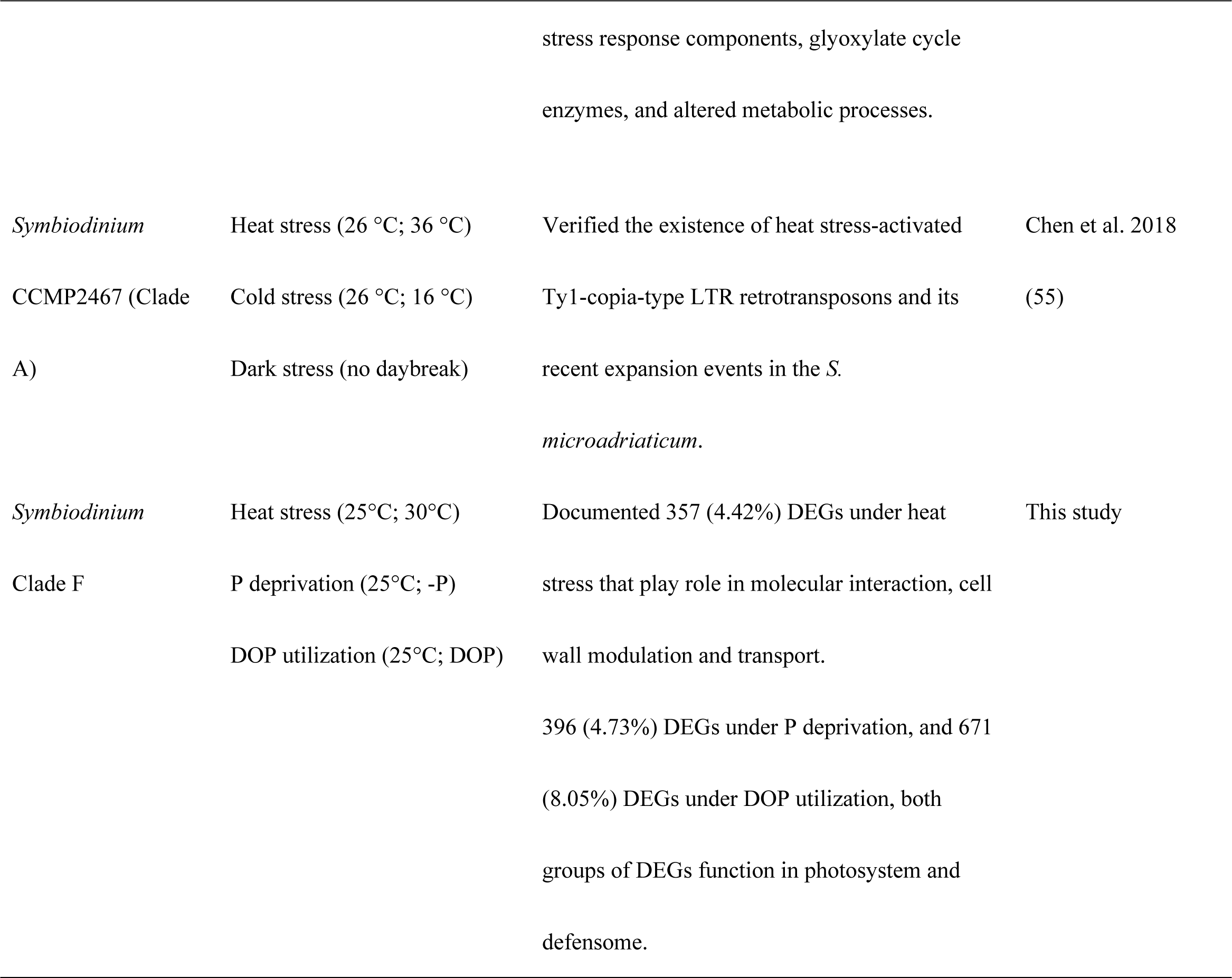
Major findings of previous heat stress related studies on *Symbiodinium*.

### “Core” genes and responsive gene groups in *S. kawagutii*

A total of 221 genes exhibited similar expression levels among the treatments, thus considered constitutive gene repertoire of *S. kawagutii* (S1 Table). We propose that this belongs to the core gene set of this species. Most of these genes function in catalytic activities and binding (Fig. 1), which are essential for metabolism and growth in the organism. To date, only a few stably expressed housekeeping genes have been identified in *Symbiodinium*, intended as reference genes with which to normalize gene expression (56, 57). The 221 stable and highly expressed core genes identified in this study add more candidate reference genes for gene expression studies on *S. kawagutii*. However, the complete core gene set very likely consist of many additional genes that were not detected in this study due to their low expression levels and our limited sequencing depth; therefore, the initial core gene set reported here will be a primer of a broader search for core genes in this and other *Symbiodinium* species in the future.

In contrast, 1,091 genes were found uniquely expressed under one growth condition or significantly differentially expressed between conditions (S2-S4 Tables), which we postulate as environmentally responsive genes (ERGs). Even though some of these “uniquely expressed” genes may also be expressed (at low levels) under other conditions examined in this study, our filtering criteria were set to reduce the likelihood. If any of these would also be expressed under other conditions not investigated in this study, it remains to be found out in the future. Of these putative ERGs, there were eight duplicated-gene families, five of which were among the expanded gene families previously identified in the genome (23). These eight showed changes in the expression level in response to all the three treatments used in this study. These are likely stress responsive genes (SRGs) in *S. kawagutii*, and potential biomarkers of stress for this species. These SRG gene families encode PPR (pentatricopeptide repeat), AR (ankyrin repeat), F-box and FNIP repeat containing proteins, EF-hand calcium-binding protein, ZnF (Zinc finger) proteins, RCC1 (regulator of chromosome condensation), glycoside hydrolase and glycosyl transferase, the functions of which will be discussed further in the following sections.

PPR proteins are organelle RNA-binding proteins to mediate gene expression at the post-transcriptional level (58, 59). The AR domain typically mediates numerous protein-protein interactions (60), and contributes to various cellular functions such as cell-cell signaling, cell–cycle regulation, and transport (61). F-box and FNIP repeat-containing proteins and AR repeat proteins have been reported to play roles in degrading proteins through protein–protein interactions in thermal sensitive type C1 *Symbiodinium* (62). EF-hand calcium binding proteins influence many Ca2^+^-dependent cellular processes (63, 64), such as cytoplasmic Ca2^+^ buffering, signal transduction protein phosphorylation, and enzyme activities (65, 66). Similarly, the diverse ZnF proteins serve as interactors of DNA, RNA, proteins and small molecules (67). GTP binding proteins of RCC1 are involved in cell cycle control and cellular processes (68, 69), and provide a possible molecular basis for permanently condensed chromatin in dinoflagellates (70, 71). Glycoside hydrolases is known to affect cell wall architecture (72, 73); therefore, its up-regulation under stress suggests a role in stress adaptation. Glycosyl transferases catalyze the transfer of sugar residues (74). Generally, it seems that these SRGs mostly function through molecular interactions, probably rendering *S. kawagutii* better adapted to environmental changes and stresses.

### Genes and encoded functions responsive to heat stress in *S. kawagutii*

Under heat stress, the most remarkable transcriptomic response included the up-regulation of a zinc finger protein (ZnF, Skav215618) and down-regulation of an ABC transporter G family member (Skav232797). ZnF in *Arabidopsis* is a positive regulator to improve thermal stress tolerance (75). The ABCG transporters have been reported to be involved in biotic and / or abiotic stress responses (76–80). The well-studied heat stress responsive genes coding for heat shock proteins (81–83) and glutathione S-transferase (84) were also up-regulated, while genes encoding conjugative proteins were down-regulated under thermal stress. Heat shock protein gene up-regulation, however, was also observed under P stress, indicating that these are likely broad stress response, rather than specific heat stress response genes. Although heat stress has been implicated more in PSII damage (85, 86), our finding of down-regulation of PSI reaction center subunit IV suggests that PSI in *S. kawagutii* is also susceptible to heat stress. Heat stress-induced elevated expression of nutrient (formate/nitrite transporter, amino acid, ion transport protein and p-type ATPase transporter) transporters suggests a higher nutrient demand and energy consumption under heat stress. In addition, two genes encoding plasma membrane permeases for high-affinity iron uptake (FTR1) were depressed under heat stress, suggesting decreased iron uptake. Meanwhile, iron binding activity was potentially promoted as inositol oxygenases and cytochrome c/P450 genes were significantly up-regulated, suggesting an increased iron demand under heat stress. *S. kawagutii* has been shown to have a higher iron requirement (500 pM soluble Fe for maximum growth rate) than other algae, although it can maintain growth at low iron availability when other trace metals such as manganese, copper or zinc is available (87). All these in concert suggest that under heat stress a fast transport system for iron (low-affinity) was functionally replacing the high-affinity iron transporter to meet the elevated demand for iron.

Our results also showed that genes encoding choline transporter-like protein 1 (CTL1), which regulates secretory trafficking of auxin (a plant hormone) transporters to control seeding growth in *Arabidopsis* (88), were down-regulated under heat stress. Meanwhile, heat stress caused up-regulation of cytokinin riboside 5’-monophosphate phosphoribohydrolase, an enzyme activating cytokinin, another plant hormone that regulates cell division and differentiation. Whether the up-regulation of this cytokinin activating enzyme gene and down-regulation of CTL1 gene would drive the cell cycle into arrest or promote cell division needs to be further investigated. Furthermore, a highly expressed gene encoding protocadherin fat (Ft) (FPKM=221-854) was strongly up-regulated under thermal stress (log2[FC]=2). In *Drosophila*, Ft, an atypical cadherin, regulates Hippo pathway and plays a key role in regulating the organ size (89, 90). Also, cadherin is responsible for cell adhesion affected by cell Ca^2+^ homeostasis, and its up-regulation is supposed to prevent apoptosis in endosymbiosis (91). It would be of interest to further explore the relationship between changes in cell size and adhesion as potential adaptive mechanism to environmental stresses and Ft expression in *Symbiodinium*. The detection of five up-regulated glycoside hydrolase genes provides evidence of cell wall change under thermal stress because these genes are potentially involved in modulating cell wall architecture (72, 73). Taken together, the results discussed above suggest that *S. kawagutii* copes with heat stress through gene regulation on cell features, including cell cycle, adhesion, cell wall architecture and physiological changes. Compared to previous findings (Table 3), the results reported here suggest that *S. kawagutii* uses unique mechanisms to cope with heat stress. Consistent with some previous studies (46), we found that photosystem II repair protein D1 gene was not among the up-regulated gene found under heat stress. It is tempting to speculate that the unique transcriptomic response (e.g. apparently elevated demand for iron, nutrients, and oxygen) may confer thermal tolerance but this requires further investigation comparing *S. kawagutii* with known heat susceptible and resistant strains on multiple physiological as well as molecular parameters.

### Genes and encoded functions responsive to P deprivation in *S. kawagutii*

Under P deprivation, the most remarkable transcriptomic response included up-regulation of reticulocyte-binding protein 2 (Skav201252) and down-regulation of photosystem I reaction center subunit IV (Skav209900). Reticulocyte-binding protein 2 is known to be involved in reticulocyte adhesion (cell-cell adhesion) in the parasite *Plasmodium falciparum* (92). Its implication in *Symbiodinium* interaction with its host cell and response to P deprivation is worth further exploring in the future. The up-regulation of alkaline phosphatase and down-regulation of metal-dependent phosphohydrolase suggest opposite functions of these enzymes in P metabolism. Alkaline phosphatase is widely known as inducible by P stress in phytoplankton to facilitate utilization of phosphomonoester type of DOP (93). The function of metal-dependent phosphohydrolase is less understood. In the dinoflagellate *Prorocentrum donghaiense*, metal-dependent phosphohydrolase protein has been shown to be differentially expressed between the cell cycle phases using quantitative proteomic analysis (94), suggesting a role of the enzyme in phosphorylation-desphosphorylation of cell cycle regulating proteins. In the roots of the land plant model *Arabidopsis thaliana* it has been shown to be down-regulated after iron deprivation (95).

Photosynthetic capacity decreases as P deficiency stress increases, as demonstrated in plants (96). Perhaps as a negative feedback to the decrease in photosynthetic capacity in *S. kawagutii*, the most abundant DEGs under P deprivation occurred in the photosystem. Among them, proteins of ferredoxin (up) and flavodoxin (down), electron transporters, showed opposite regulation under P deprivation (Fig. 4). They can replace each other in the photosynthetic electron transfer chain of cyanobacteria and algae (97). In these photosynthetic taxa, flavodoxin is induced by iron deficiency while the iron-containing ferredoxin is down-regulated (28–31). The up-regulation of haem-binding GO terms observed under P deprivation suggests a higher demand for iron under P stress, similar to the case of heat stress. Thus, the up-regulation of flavodoxin might be an adaptive response to alleviate iron stress. DEGs involved in the production of chemical energy were all up-regulated, suggesting energy deficiency under P stress. DEGs enriched in oxidoreductase activity and heme binding were probably involved in P stress. The increased expression of abiotic defense genes (efflux, oxdative and antioxidant protein genes, and heat shock proteins) suggests these genes are also P stress responsive.

Furthermore, we observed suppressed expression of two variants of the meiosis associated gene mei2 under P deprivation. MEI2 is a RNA-binding protein involved in meiosis, crucial for commitment to meiosis (i.e. switching from mitotic to meiotic cell cycle) in the fission yeast *Schizosaccharomyces pomb* in which this gene is known to be induced by N-nutrient starvation (98–100). The opposing response of this gene to N (in yeast) and P deprivation (in *Symbiodinium*) is interesting, suggesting that P deficiency might inhibit meiosis whereas N starvation may induce it leading to encystment. These might indicate that under P stress, *S. kawagutii* meiosis was repressed, which should a topic of interest for future research.

### Genes and encoded functions responsive to DOP replacement in *S. kawagutii*

When grown on Gro3P as the sole source of P, the most remarkable transcriptomic response in *S. kawagutii* included up-regulation of reticulocyte-binding protein (Skav201252) and down-regulation of ATP-grasp fold (Skav221360) compared to the control (grown on phosphate). As it was also up-regulated under P deprivation, reticulocyte-binding protein up-regulation seems to be responsive to a change of P condition. The ATP-grasp hold is one of the ATP-grasp superfamily, which includes 17 groups of enzymes, catalyzing ATP-dependent ligation of a carboxylate containing molecule to an amino- or thiol-containing molecules (101), contributing to macromolecular synthesis. Its down-regulation under Gro3P treatment suggests reduction of macromolecular synthesis when glycerophosphate is supplied as the sole P-source.

Compared to DIP growth condition, acid phosphatases were uniquely up-regulated under Gro3P, suggesting its potential roles in DOP utilization. Furthermore, similar defensome sets as found under P deprivation were also observed under DOP, which in addition also induced ABC transporter and major facilitator superfamily. The two membrane transporters couple solute movement to a source of energy (102), which potentially plays a vital role in transferring Gro3P. DnaJs play important roles in protein translation, folding, unfolding, translocation, and degradation, and regulates the activity of Hsp70s (103). The abundant and up-regulated DnaJ as well as Hsp70 (Fig. 4) were both induced by DOP condition in our study. The roles of Hsp genes in DOP metabolism remains to be further explored.

The largest enriched GO term under DOP condition was integral component of membrane, suggesting high transport activities to utilize DOP. It is interesting to note that auxin efflux carrier was induced by organophosphate replacement but not by P limitation. Auxin is a plant hormone, previously shown to occur and influence development in algae (104). Auxin efflux carrier proteins influence many processes in plants including the establishment of embryonic polarity, plant growth, apical hook formation in seedlings and the photo- and gravitrophic responses (105–107). In rice, the auxin efflux carrier gene is involved in the drought stress response (108). In the genome of *S. kawagutii*, besides the 4 auxin efflux carrier genes, there are three auxin responsive GH3 genes, indicative of an auxin-based gene regulatory pathway in this species. It is unclear what physiological consequence the elevated expression of the auxin efflux carrier gene would lead to, but potentially it may be responsible for promoting cellular growth under DOP condition.

Utilization of Gro3P also induced differential gene expression related to carbohydrate metabolic process and tricarboxylic acid cycle. Glyceraldehyde 3-phosphate dehydrogenase (GAPDH) and glycerol-3-phosphate dehydrogenase (GPDH) genes were greatly up-regulated. GAPDH interacts with different biomolecules, and has been known to play an important role in diatom’s ecological success (109). In Chironomidae, GAPDH enhances heavy metal tolerance by adaptive molecular changes through binding at the active site (110). GPDH is a very important enzyme in intermediary metabolism and as a component of glycerophosphate shuttle it functions at the crossroads of glycolysis, oxidative phosphorylation and fatty acid metabolism (111). GAPDH- and GPDH-dependent metabolic pathways seem to be modulated by Gro3P utilization in *S. kawagutii*, and yet the physiological or ecological implications still remain to be further investigated.

## Conclusions

This study is the first to explore transcriptomic responses to P deprivation, DIP replacement with DOP, as well as thermal stress for the same genotype of *Symbiodinium*, a strain that has been less frequently studied. With an analysis strategy to ameliorate impact of sequencing and replicate limitation, we focused on highly expressed common (core) genes of the species and differentially expressed genes responding to specific treatments. We identified 221 (2.04%) such core genes for *S. kawagutii*, which mostly were in the Gene Ontology terms of catalytic activity and binding. Differentially gene expression showed that eight duplicated gene families were in response to all the three treatments investigated in this study, including ankyrin repeat (AR)-, F-box and FNIP repeat- and pentatricopeptide repeat (PPR)-containing proteins, and EF-hand domain-containing calcium-binding proteins, Zinc finger (ZnF) proteins, glycoside hydrolases, regulator of chromosome condensation (RCC1), and glycosyl transferases. These apparently are non-specific stress response genes in this species. Specific to heat stress, 357 (4.42%) genes were found to be differentially expressed, with predicted roles in active molecular (protein-protein/RNA/DNA) interaction, cell wall modulation and transport of iron, oxygen, and major nutrients, in addition to the expected up-regulation of heat shock protein genes. We did not observe a significant up-regulation of photosystem II repair protein D1 gene as expected under heat stress, which along with the distinct transcriptomic response observed suggests that this species has a unique mechanism by which to cope with heat stress and is possibly thermal tolerant. Our results also indicate that there is likely a higher demand for nutrients, iron, and oxygen under heat stress.

About as many DEGs (396, 4.73%) were identified under P deprivation while nearly double of that (671, 8.05%) were detected under DOP (glycerophosphate) utilization; in both cases most of the DEGs were up-regulated and predicted to function in photosystem and defensome, indicating that photosynthesis and defense are probably the most markedly impacted physiologies under varying P-nutrient conditions. This study provides novel insights into responses in *S. kawagutii* to both temperature and P nutrient variations, setting a framework for more transcriptomic research in the future on *Symbiodinium* spp. to uncover common and stress-specific features of stress sensitive (bleaching prone) and tolerant strains and to elucidate triggers of coral bleaching.

## Materials and Methods

### *S. kawagutii* cultures, sampling and RNA sequencing

*S. kawagutii* strain CCMP2468 was obtained from the National Center for Marine Algae and Microbiota (NCMA) in the Bigelow Laboratory of Ocean Science (Boothbay Harbor, Maine, USA). It was maintained in L1 medium prepared from natural seawater under an illumination of ~200 μE m^-2^ s^-1^ with a 14:10 light dark cycle and at a temperature of 25°C. For the experiment, 1 L of the cells in the exponential growth stage were divided into 4 bottles and collected by centrifugation at 3,000 g, 25°C for 15 min to remove the old cultural medium, and re-suspended with different growth media and divided into 4 groups (SymkaSL1-4), each in triplicate. SymkaSL1 and SymkaSL2 were grown in artificial seawater enriched with normal standard L1 medium (nutrient complete), SymkaSL3 in artificial seawater-based L1 medium with deprived P (P-), and SymkaSL4 in artificial seawater-based L1 medium with DIP replaced by glycerol-3-phosphate (Gro3P) at equivalent concentration (36.2 μM). Each subsample was washed with the corresponding cultural medium 3 times to remove trace old culture medium and transferred to triplicated 150-mL bottles. SymkaSL1 bottles were transferred to 30 °C for thermal stress treatment, while SymkaSL2-4 were kept at 25 °C. After one-week incubation under those conditions, cells were harvested for RNA extraction.

RNA was isolated using Trizol reagent (Life Technologies, Grand Island, NY) coupled with QIAGEN RNeasy kit (QIAGEN Inc., Germantown, MD) according to Zhang et al. (112). The RNA samples were subjected to RNA-seq in 2 x 50bp paired-end format at the National Center for Genome Resources under Marine Microbial Eukaryote Transcriptome Sequencing Project (MMETSP) (35). Before sequencing, RNA extracts from the triplicate cultures were pooled at equal RNA quantity (due to high cost of sequencing back then). Raw data was uploaded to NCBI under the accession numbers SRR1300302, SRR1300303, SRR1300304 and SRR1300305.

### Data preprocessing

The genome reference datasets of *S. kawagutii* were downloaded from http://web.malab.cn/symka_new/index.jsp for transcriptomic reads mapping. Cutadapt (113) was used to remove adaptor sequences with parameters of “-e 0.05 –overlap 25 –discard-trimmed -m 20” for each end of fastq data separately. The trimmed clean reads were evaluated with FastQC to check quality. Multiqc software was used to integrate FastQC reports.

### Reads counting

Clean reads were aligned to the genome sequences by HISAT2 (114) with default parameters. The generated sequence alignment map (SAM) was then converted into its binary format, BAM, and sorted using SAMtools (115). Mapping quality was evaluated using Qualimap (bamqc) (116) based on BAM file. Read summarization was analyzed with featureCounts, and multi-mapping reads were not counted (parameters of -p -C). Read counts were normalized as <u>F</u>ragments <u>P</u>er <u>K</u>ilo-base of exon model per <u>M</u>illion mapped fragments (FPKM). For each pair of comparison, to avoid inflation of differential expression by low-expression genes, genes with read counts per million (CPM) < 10 in both samples, or not detected in one sample while CPM < 20 in the other sample, were excluded. The remaining genes (herein named actively expressed genes or AEG) were subjected to further statistics analyses.

### Identification of expressed core genes

Commonly as well as highly expressed genes in *S. kawagutii* grown in all four conditions were identified as core genes. Several steps were taken. Firstly, from the AEG, genes commonly detected in four samples were identified (AEG-C). Secondly, for each gene in SGS-C, the average FPKM and coefficient of variance (CV) in the four samples were calculated. Finally, Genes showing expression of ≥ 50th percentile average FPKM and low CV of ≤ 0.1 were collected and named cores genes.

### Differential gene expression analysis

To generate reliable differential gene expression profiles in the absence of biological triplicates (due to the pooling of the triplicate culture samples), two methods, edgeR and NOIseq in R, were used in parallel and only the consistent results from both methods were used for further analysis. Both methods account for biological variability when samples have no replicates. edgeR determines DEGs using empirical Bayes estimation as well as exact tests based on negative binomial models (117), and is widely used for analyzing DEGs for non-replicated samples. The moderate biological coefficient of variation (BCV) of 0.2 was used to estimate dispersion. DEGs of edgeR were screened with threshold of FDR ≤0.05 and the absolute value of log2Ratio ≥1. NOIseq is a non-parametric approach consisting of NOISeq-real and NOISeq-sim. NOISeq-sim simulation of noise distribution in the absence of replication was optimized. The parameters of q=0.8, pnr=0.2, nss=5 and v=0.02 were set as previously suggested (37). Only genes identified as DEGs by both methods were classified as true DEGs. Heatmap was used to display DEGs with value of log2(RPKM+1) transformation.

### Gene ontology and KEGG functional enrichment

To optimize Gene Ontology (GO) annotation rate, InterProscan tool was used to scan protein signatures of the newest updated databases (version 5.27-66.0, lookup_service_5.27-66.0) against *S. kawagutii* genome protein sequences. GO annotation results were visualized by WEGO 2.0 (118). KO information was grabbed from genome Kyoto Encyclopedia of Genes and Genomes (KEGG) annotation information. We used significant DEGs as a foreground to perform GO or KEGG functional enrichment analyses with ClusterProfiler package (119). And enrichment visualization was displayed with enrichplot package (120). Because of largely unexplored genome of *Symbiodinium* spp., annotation rate of GO and KEGG were generally low.

## Acknowledgments

We thank Tangcheng Li and Xiaohong Yang for their technical assistance. Thanks are also due to Mr. Jon Kaye, Dr. Stephanie Guida and the rest of the MMETSP team throughout the project for their generous support in various capacities with the sequencing program and the initial processing of the sequencing output.

## Supporting information

**S1 Table. Information of core genes commonly expressed under all the four culture conditions (heat stress, control, P-depleted and DOP replacement) in *S. kawagutii***. (XLSX)

**S2 Table. Information of differentially expressed genes in response to heat stress in *S. kawagutii***. (XLSX)

**S3 Table. Information of differentially expressed genes in response to phosphorus stress in *S. kawagutii***. (XLSX)

**S4 Table. Information of differentially expressed genes in response to dissolved organic phosphorus (DOP, glycerophosphate) replacement in *S. kawagutii***. (XLSX)

